# IMPROVING ALPHAFOLD2 PERFORMANCE WITH A GLOBAL METAGENOMIC & BIOLOGICAL DATA SUPPLY CHAIN

**DOI:** 10.1101/2024.03.06.583325

**Authors:** Geraldene Munsamy, Tanggis Bohnuud, Philipp Lorenz

## Abstract

Scaling laws suggest that more than a trillion species inhabit our planet but only a miniscule and unrepresentative fraction (less than 0.00001%) have been studied or sequenced to date. Deep learning models, including those applied to tasks in the life sciences, depend on the quality and size of training or reference datasets. Given the large knowledge gap we experience when it comes to life on earth, we present a data-centric approach to improving deep learning models in Biology: We built partnerships with nature parks and biodiversity stakeholders across 5 continents covering 50% of global biomes, establishing a global metagenomics and biological data supply chain. With higher protein sequence diversity captured in this dataset compared to existing public data, we apply this data advantage to the protein folding problem by MSA supplementation during inference of AlphaFold2. Our model, BaseFold, exceeds traditional AlphaFold2 performance across targets from the CASP15 and CAMEO, 60% of which show improved pLDDT scores and RMSD values being reduced by up to 80%. On top of this, the improved quality of the predicted structures can yield better docking results. By sharing benefits with the stakeholders this data originates from, we present a way of simultaneously improving deep learning models for biology and incentivising protection of our planet’s biodiversity.

## 1 Introduction

In the last several years we have experienced the rise of a plethora of deep learning models applied to a wide range of biological tasks [1], [2]. Of particular prominence is the protein folding problem, given its impact on structural biology and drug discovery, for which AlphaFold2, RoseTTAFold, and ESMFold, for example, are providing promising and often highly accurate predictions [3], [4], [5]. A lot of research and effort has been put into optimising the architecture of these models to improve performance [6]. However, given that these models depend on the protein sequence and structure datasets available for training, we deployed a data-centric approach towards improving deep learning models in biology, exemplified on the protein folding problem for the purpose of this study.

Previous studies have shown that with improved data quality and quantity, the error loss of transformers while training would no longer follow a power law, but rather an exponential relationship [7]. When looking at the public sequence databases available for deep learning in biology, such as UniProt, NCBI, or MGnify, scaling laws suggest that these datasets represent less than 0.000001% of life on earth [8], [9], [10]. A significant portion of sequences deposited in these databases originate from human, mammals, and model organisms that are cultivated in narrow laboratory conditions [11], [12]. Furthermore, for environmentally collected sequence data, these resources lack consistent geolocation and environmental metadata. The latter point not only precludes us from inferring how comprehensively life on earth is represented in these databases, but also raises questions relating to the governance of biological sequence data.

In the case of environmentally collected biological samples for genomic (or other -omic) purposes, ethical compliance to Access and Benefit Sharing (ABS) agreements is contingent on both explicit prior informed consent (PIC) and mutually agreed terms (MAT) that speak to the potential for commercialization, and consistent traceability from a sequence to its sampling origin. Regulatory frameworks ensuring ethical ABS upon commercialization of biological resources have been driven on an international level by the United Nations Convention on Biological Diversity (CBD) and are documented in the 2011 Nagoya Protocol [13]. The inclusion of digital sequence information as part of ABS frameworks is an area of significant development in this context [14]. Historically, there are many instances where materials analyzed for research purposes have contributed to commercial assets of unexpectedly high value without due reconsideration of the original agreements under which the samples were accessed and what fair benefit sharing should look like, leading to controversies around biopiracy and impediments to the development of assets that could have been transformative for industry and human health [15], [16], [17], [18].

Here we describe a global metagenomics and biological data supply chain that simultaneously addresses both the issue of equitable benefit sharing of digital sequence information and the large knowledge gap we experience regarding genomic sequence diversity of life on earth with the aim to improve biological deep learning models. The genome and protein sequences as well as consistently collected metadata derived from metagenomic sampling expeditions are captured in a knowledge graph that counts 6 billion relationships at the time of writing. In the context of knowing that models like AlphaFold2 perform less well on orphan proteins for which deep multiple sequence alignments (MSAs) cannot be generated [19], we show that the performance of AlphaFold2 can be improved when MSAs are supplemented with diverse sequences from our knowledge graph. Assessing confidence and accuracy of the predicted structure, we observe the root mean squared deviation (RMSD) compared to ground-truth crystal structures being reduced by up to 80% . We display improved structure predictions for a wide range of CASP15 and CAMEO competition targets [20], [21], and demonstrate that docking performance can be improved as a result, too.

## 2 A global metagenomic and biological data supply chain addresses the knowledge gap of biological sequence diversity

In order to curate genomic and biological data that are more representative of the true diversity of life on earth, we entered Access Benefit Sharing (ABS) agreements following prior informed consent (PIC) of relevant landowners and stakeholders across 23 nations on 5 continents before conducting environmental metagenomic sampling alongside geological, geographic, and chemical metadata collection (Figure 1 A). The sampling sites cover 50% of global biomes according to the WWF Ecoregion defintion [22]. Methods pertaining to the sampling, sequencing, and bioinformatic assembly and annotation following these expeditions are described in Supplementary Section A1.

**Figure 1:**
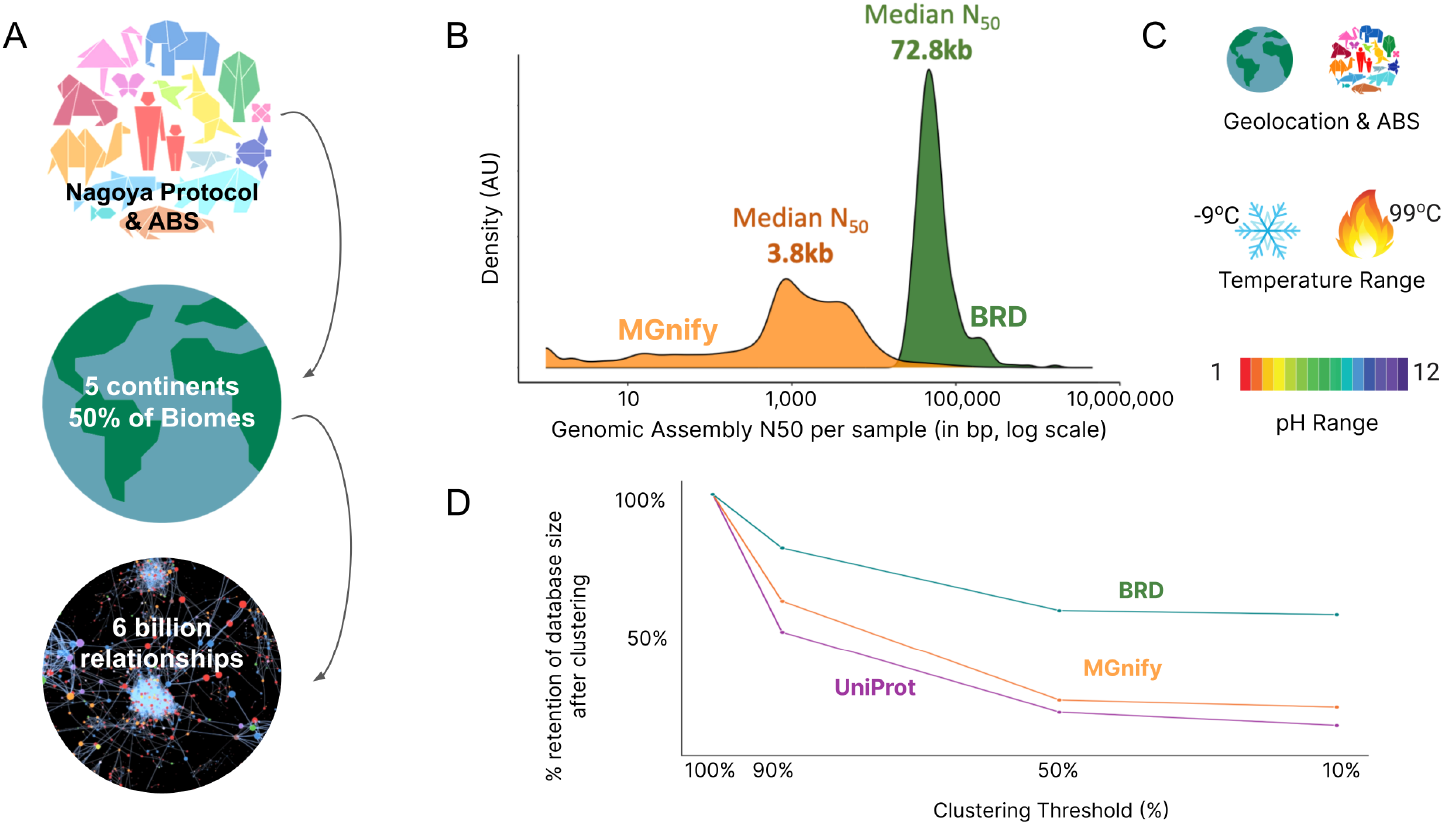
Strategy for accessing and organising data derived from a global metagenomic and bilogical data supply chain. A. Biological and metagenomic sequence collection strategy covering ABS agreements & Nagoya compliance; global expeditions covering 5 continents; and organisation of this data into a knowledge graph (data resource hereforth referred to as BRD). B. Metagenomic assembly length distribution as measured by the N50 value for MGnify and BRD. C. Examples of metadata and features captured in BRD that other resources lack or do not consistently display. D. Protein sequence diversity of MGnify, UniProt, and BRD, as shown by clustering the sequence content.

We organised all genome and protein sequences alongside chemical and environmental metadata into a knowledge graph counting 6 billion relationships at the time of writing. With the downstream application of MSA generation for AlphaFold2 predictions in mind, we wanted to ensure that the sequences for such application are derived from high-quality and long (meta)genomic assemblies. The reasoning for this is that a large portion of the sequence database content that AlphaFold2 currently derives MSAs from is MGnify, and a significant portion of the metagenomic assemblies found in MGnify are fragmented and not long enough to cover entire open-reading frames (ORF) for larger proteins [10]. The length distributions of metagenomic assemblies in MGnify and our database (from hereon referred to as BRD, Basecamp Research Data) are displayed in Figure 1B. With consistent metadata collection we were able to sample a wide range of geological and chemical environments, spanning a temperature range of -9 to 99°C (15.8 to 210°F), and a pH range of 1 to 12, as shown in Figure 1C. We then assess the diversity of the protein sequences deposited in BRD compared to MGnify and UniProt by comparing how the size of the databases collapses when clustered at 90%, 50%, and 10% (Figure 1D).

## 3 Improving AlphaFold2 through MSA augmentation

To leverage the sequence diversity captured in BRD for MSA supplementation during inference without sacrificing too much speed for sequence search, we clustered both MGnify and BRD at a 50% identity threshold using MMseqs2 Linclust [23]. The resulting combined metagenomic sequence dataset contained approximately 1 billion sequences. To assess whether the addition of sequences through MSA supplementation would improve AlphaFold2, we performed structural analysis on sequences from the CASP15 and CAMEO targets.

### 3.1 CASP15 targets

CASP (Critical Assessment of Structure Prediction) is a biennial global experiment designed to advance the state of the art in modeling the three-dimensional structure of a protein from its amino acid sequence. Organized by the scientific community, it invites participants to present their modeling predictions for a selection of proteins whose experimental structures are yet to be deposited in the Protein Data Bank (PDB) [24].

We predicted the structures of 49 CASP15 regular targets, among which single monomeric protein crystal structures were available in the PDB, establishing a benchmark for our comparative study. The structural predictions were evaluated using the predicted Local Distance Difference Test (pLDDT) scores, providing a per-residue confidence metric ranging from 10 to 100 [25]. Among the 49 targets analyzed, 61.22% demonstrated an improvement in pLDDT scores, with increases ranging from 0.08 to 24. The scores of these targets are provided in Supplementary Information Table 1. For the subset of targets where we did not observe an increase in pLDDT the average percentage difference was 3.1%.

**Table 1:** CASP15 targets that display an increase in the pLDDT scores for BaseFold predictions compared to AlphaFold2.

| ID | AlphaFold2 | BaseFold | Difference |
| --- | --- | --- | --- |
| T1176 | 69.94 | 94.61 | 24.67 |
| T1113 | 69.83 | 90.64 | 20.81 |
| T1147 | 73.50 | 92.15 | 18.64 |
| T1114s1 | 62.72 | 78.22 | 15.50 |
| T1137s7 | 71.48 | 82.15 | 10.68 |
| T1134s2 | 77.90 | 88.43 | 10.53 |
| T1106s1 | 69.24 | 79.41 | 10.17 |
| T1115 | 75.16 | 85.02 | 9.85 |
| T1119 | 66.97 | 72.45 | 5.48 |
| T1170 | 90.57 | 94.24 | 3.67 |
| T1137s8 | 83.33 | 86.92 | 3.59 |
| T1124 | 88.89 | 91.13 | 2.24 |
| T1137s9 | 84.69 | 86.89 | 2.19 |
| T1139 | 89.52 | 91.40 | 1.88 |
| T1155 | 79.62 | 81.43 | 1.81 |
| T1150 | 92.90 | 94.57 | 1.67 |
| T1114s2 | 86.56 | 88.11 | 1.55 |
| T1109 | 94.48 | 95.94 | 1.46 |
| T1153 | 86.36 | 87.82 | 1.46 |
| T1133 | 83.97 | 85.23 | 1.27 |
| T1120 | 92.24 | 93.45 | 1.21 |
| T1157s2 | 89.17 | 90.29 | 1.12 |
| T1110 | 94.87 | 95.95 | 1.08 |
| T1106s2 | 94.48 | 95.30 | 0.82 |
| T1134s1 | 95.08 | 95.88 | 0.81 |
| T1145 | 95.07 | 95.83 | 0.76 |
| T1158 | 82.79 | 83.43 | 0.64 |
| T1127 | 93.01 | 93.28 | 0.27 |
| T1157s1 | 84.40 | 84.49 | 0.09 |

**Table 2:** Sequence contributions by MGnify and BRD to the MSAs of the following CASP15 and CAMEO targets.

| Target | Mgnify | BRD |
| --- | --- | --- |
| T1158 | 163 | 337 |
| T1145 | 426 | 74 |
| T1134s1 | 307 | 193 |
| T1106s2 | 163 | 121 |
| T1133 | 101 | 399 |
| T1124 | 149 | 351 |
| T1137s8 | 108 | 397 |
| T1106s1 | 19 | 12 |
| T1137s2 | 44 | 456 |
| T1147 | 104 | 396 |
| T1113 | 1 | 7 |
| T1176 | 12 | 3 |
| 8HWI | 291 | 209 |
| 8SSD | 174 | 326 |
| 8U00 | 105 | 395 |
| 8JYT | 80 | 420 |
| 8DYD | 404 | 96 |

Subsequently, we calculated the Root Mean Square Deviation (RMSD), which quantifies the mean distance between corresponding atoms of superimposed protein structures [26]. RMSD is a critical metric in CASP competitions for gauging the congruence of predicted protein structures with their experimentally determined counterparts. An RMSD value ranging from 0 to 3 Å ngstroms denotes a high level of structural similarity, particularly in the protein backbone, indicative of a more accurate prediction. For all targets that had an increased pLDDT score the RMSD score was computed using the SwissModel server [27] which revealed an RMSD score reduction ranging from 0.02 to 3.33. We show an overview of targets from CASP15 where MSA supplementation both improves the pLDDT and reduces the RMSD score in Figure 2. We visualized two specific examples with structural superimposition and corresponding MSA visualization as phylogenetic trees in Figure 3A and 3B.

**Figure 2:**
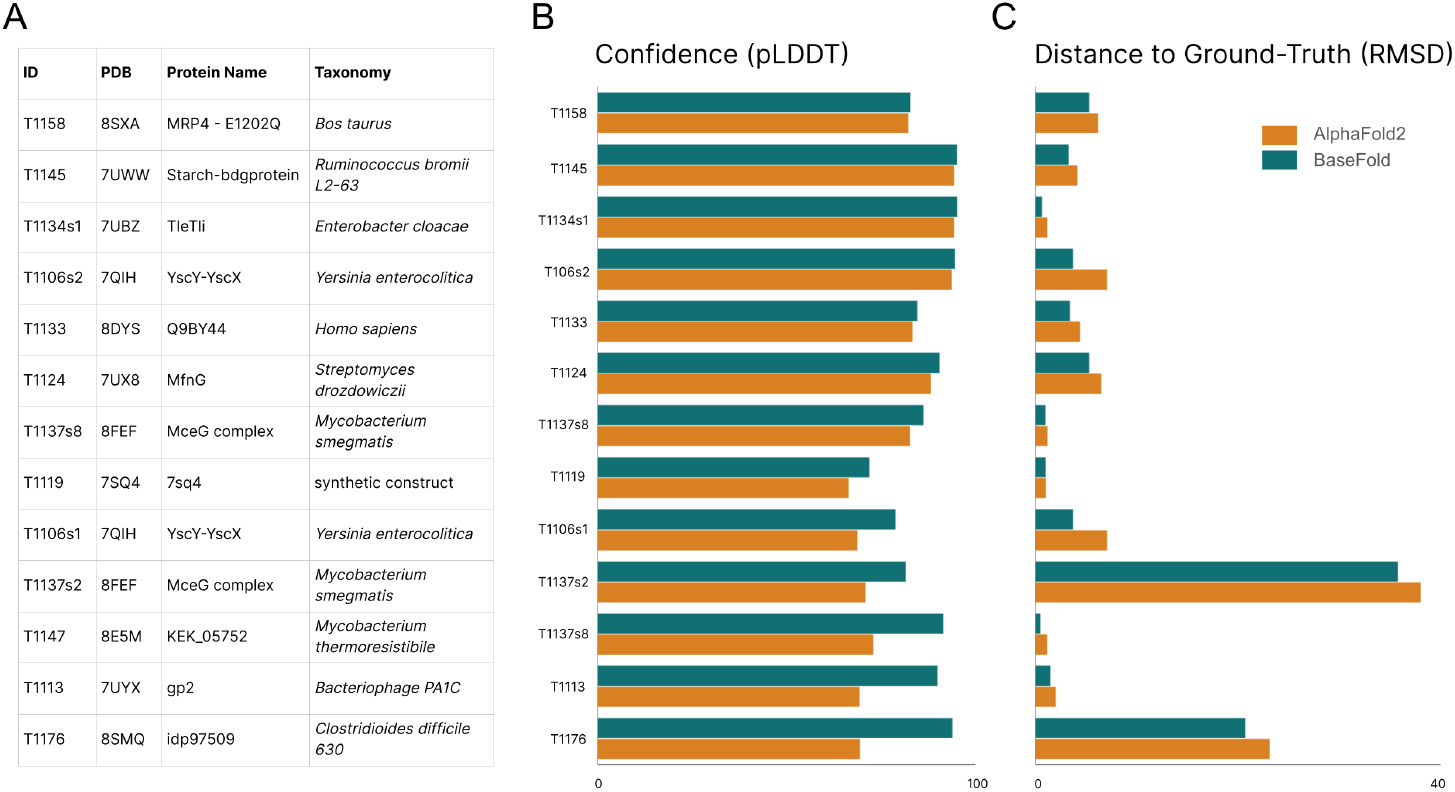
MSA supplementation improves AlphaFold2 performance across CASP15 targets. A. Table of targets where MSA supplementation improves both pLDDT score (shown in B) and RMSD scores (shown in C).

**Figure 3:**
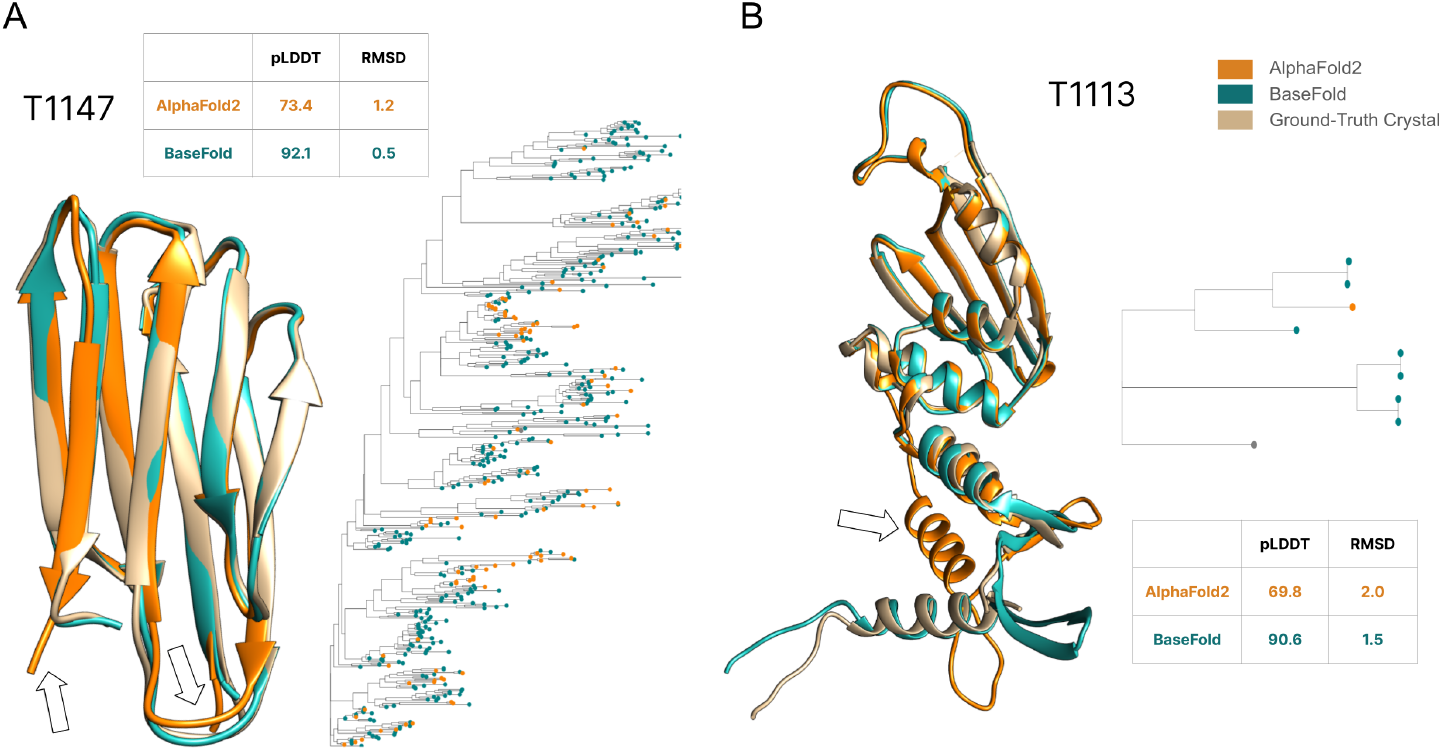
Structural superimposition and corresponding phylogenetic trees derived from MSAs for CASP15 targets T1147 (A) and T1131 (B). Significant discrepancies between the AlphaFold2 prediction and the crystal structure are indicated with a white arrow.

### 3.2 CAMEO targets

Continuous Automated Model Evaluation (CAMEO) is an online platform that offers automated assessments of 3D protein prediction models, providing weekly updates based on sequences awaiting deposition in the PDB[28]. Expanding to address the structural bioinformatics community’s evolving needs, CAMEO features a variety of assessment categories, including prediction coverage, local accuracy, and completeness, while maintaining a focus on evaluating quality estimates for protein structure predictions.

In this study, we predicted the structures of 26 CAMEO targets, predominantly comprising medium to hard difficulty levels. Notably, 57% of these targets demonstrated an increase in pLDDT scores ranging from 0.03 to 10.04. We show an overview of targets from CAMEO where MSA supplementation both improves the pLDDT and reduces the RMSD score in Figure 4 and visualized two specific examples with structural superimposition and corresponding MSA visualization as phylogenetic trees in Figure 5A and 5B.

**Figure 4:**
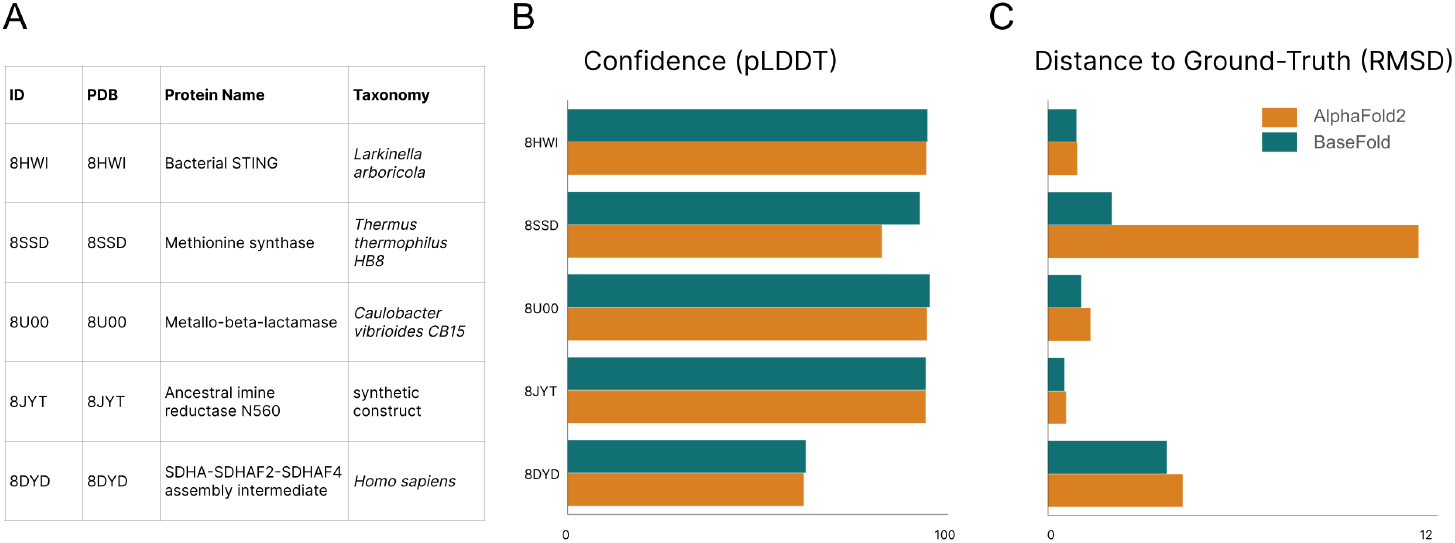
MSA supplementation improves AlphaFold2 performance across a range of CAMEO targets. A. Table of targets where MSA supplementation improves both pLDDT score (shown in B) and RMSD scores (shown in C).

**Figure 5:**
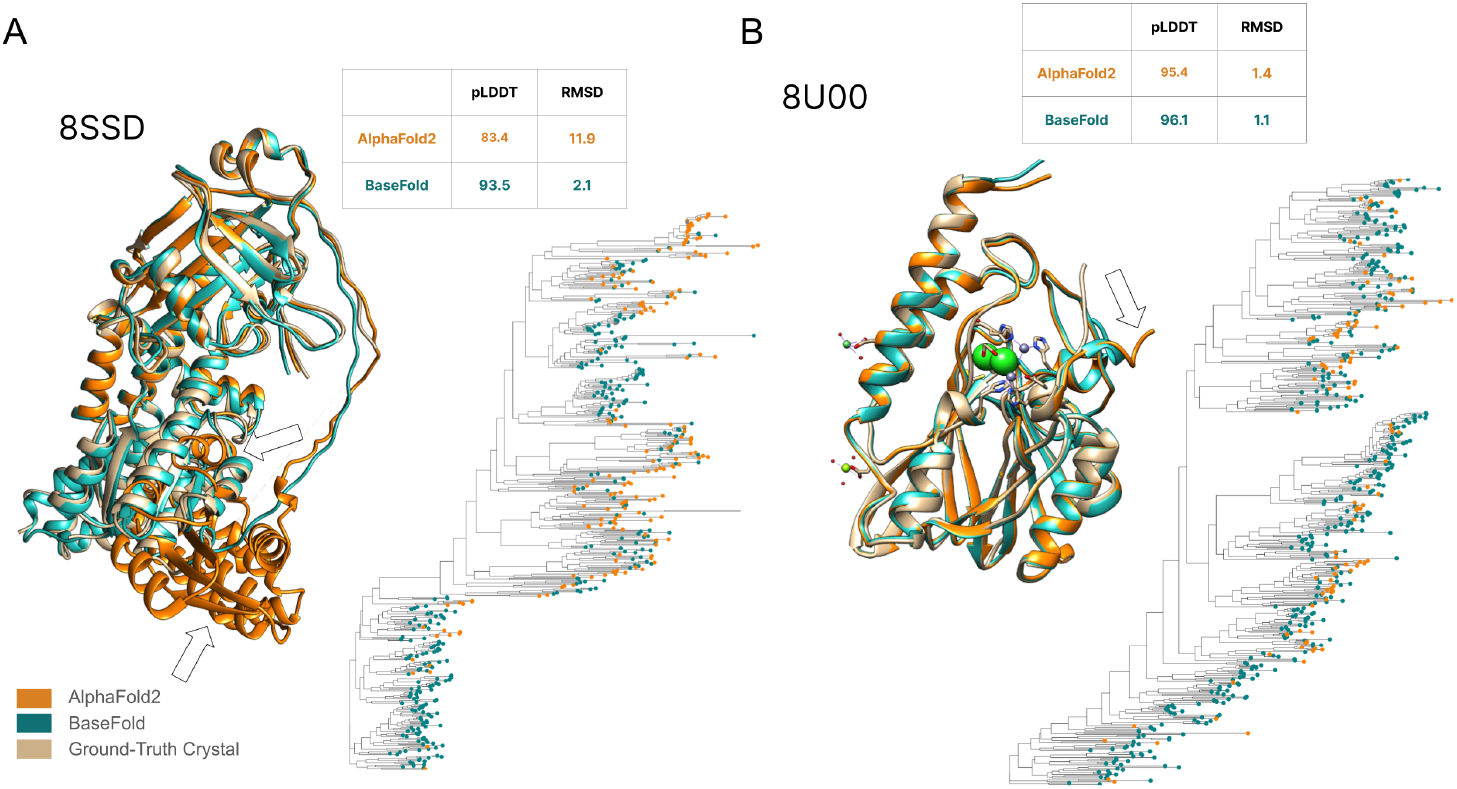
Structural superimposition and corresponding phylogenetic trees derived from MSAs for CAMEO targets 8SSD (A) and 8U00 (B). Significant discrepancies between the AlphaFold2 prediction and the crystal structure are indicated with a white arrow.

### 3.3 Improving the scale of BaseFold

Building diverse MSAs requires large compute capabilities and is time consuming. To ensure quicker iterations and greater scalability in structure predictions we refined the MSA generation step. We implemented the same strategy implemented by ColabFold [29] for the database preparation in addition to creating two environmental databases to search against which contained BRD clustered at 50% and 90% respectively. More information on the clustering of the respective BRD databases can be found in Supplementary Section A2. We ran these versions of BaseFold using the default settings and predicted the structure of 395 medium and hard CAMEO targets between 2023-02-25 to 2024-02-27 setting the template date to 2023-01-01. We visualise this in Figure 6, where we see that optimisation for speed at lower clustering thresholds does not impact performance significantly.

**Figure 6:**
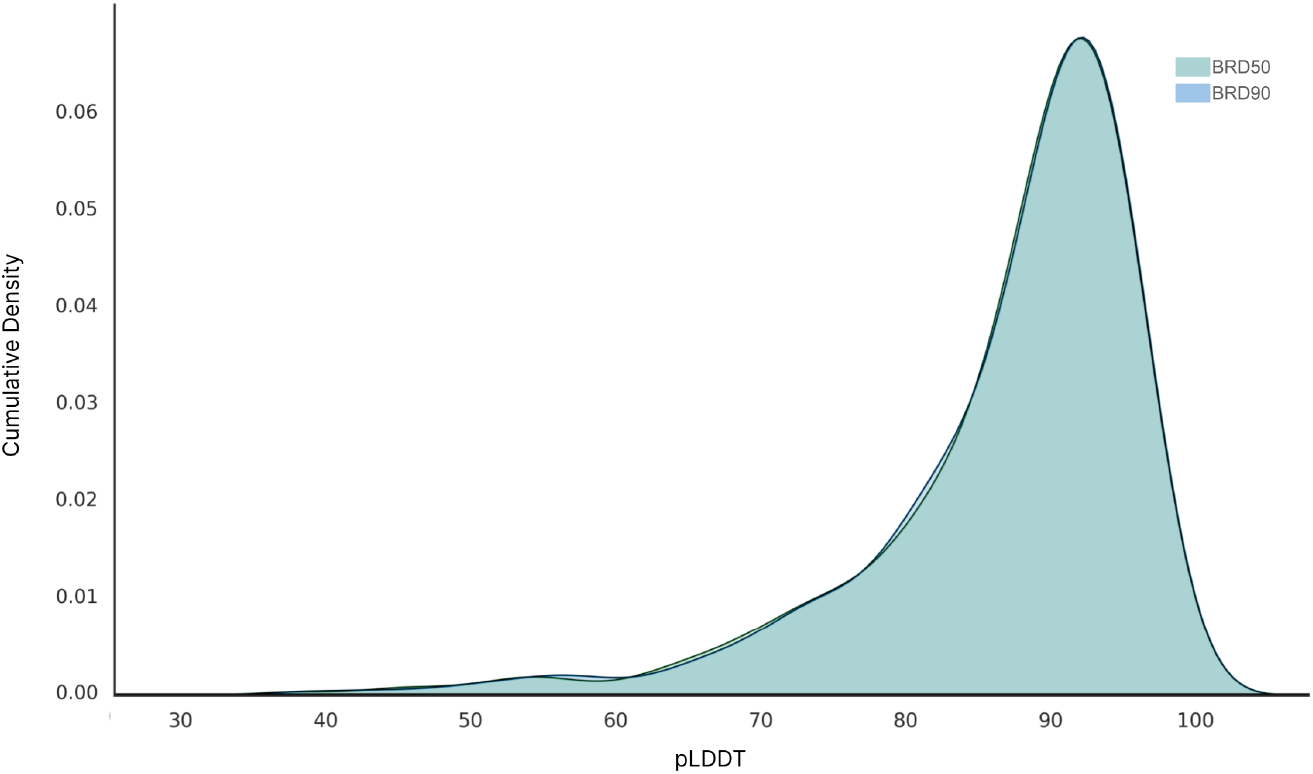
pLDDT distribution of CAMEO targets at 50% and 90% suggest that optimisation for speed at lower clustering thresholds does not impact performance significantly.

**Figure 7:**
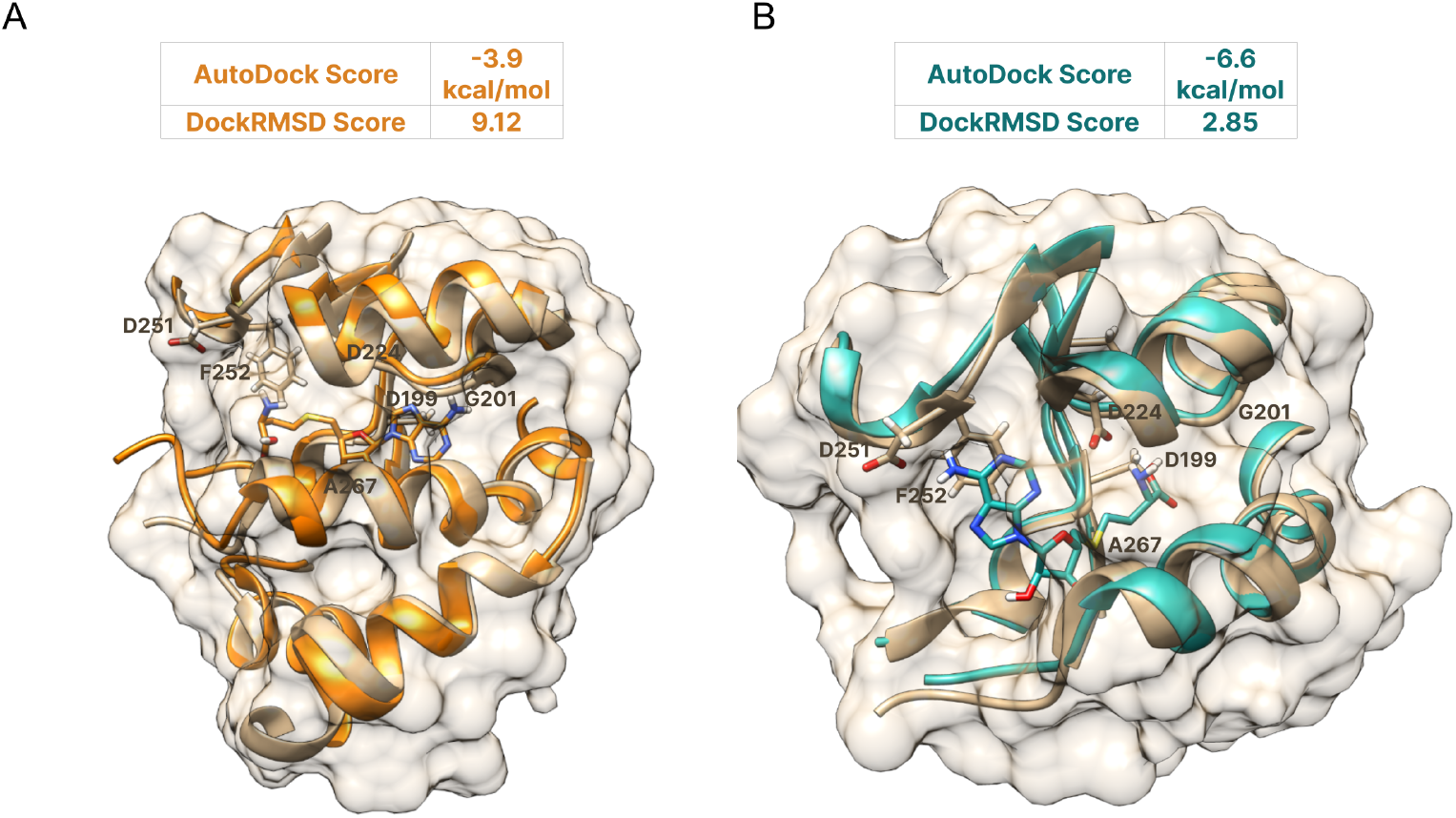
Docking of SAH to MFnG (an L- and D-tyrosine O-methyltransferase, T1124) is improved significantly by MSA-supplemented structure prediction Predicted structure, superimposed to the ground-truth crystal with docked substrates are shown for traditional AlphaFold2 (A) and the MSA-supplemented version (B). Active site residues are highlighted and docking scores indicated in respective tables.

### 3.4 Molecular Docking

Across the structure prediction improvements shown above, we noticed a particularly significant improvement for CASP15 target T1124 (an L- and D-tyrosine O-methyltransferase, MFnG). Using this example, we wanted to assess the impact of the structural prediction methods on substrate conformation and binding affinity. We performed molecular docking on the MSA-augmented and AlphaFold2-derived structure of MFnG to its substrate S-adenosyl-L-homocysteine (SAH). Docking was performed using AutoDock Vina [30], more information of the docking protocol can be found in Supplementary Section A7. The MSA-augmented structure, when bound to SAH, achieved the lowest binding score of -6.6 kcal/mol, in comparison to the AF2 structure bound to SAH, which yielded a docking score of -3.9 kcal/mol. Further analysis of the docked complexes was assessed using DockRMSD [31]. This tool measures the RMSD between two poses of the same ligand molecule docked onto the same protein structure, without presuming a known atomic order between the two files. The DockRMSD score for the MSA-augmented structure bound to SAH was 2.85 Å ngstroms, while for the AF2 structure bound to SAH, it was 9.12 Å ngstroms.

## 4 Discussion

Relative to the magnitude of diversity of life on earth – whether taxonomically or with respect to genomic or protein sequence space – everything that has been captured in public data to date still only represents a tiny fraction. By building a data supply chain in partnership with biodiversity stakeholders we aim to leverage this data to continuously improve deep learning models in biology. Specifically for protein folding, we have demonstrated that by supplementing MSAs with diverse sequences from this supply chain, we can improve AlphaFold2 predictions for a range of targets from the CASP15 and CAMEO competitions. Depending on the protein family and breakdown of the sequence composition of the corresponding MSAs generated during inference, our supplementation approach can improve AlphaFold2 predictions substantially, with some RMSD values (deviation from the ground-truth crystal) decreasing by over 80%. We show that improvements as significant as this can also improve substrate/ligand docking performance. We envision this will positively impact enzyme engineering and drug discovery efforts.

Regarding further work, we foresee additional analysis on the sequence composition of reference databases and what the ideal breakdown of sequence space should look like, in a way that balances both inference speed and accuracy of the predicted structure. Moreover, with further data collection in alignment with the United Nations’ ABS principles, we aim to continue unifying the goal of biodiversity conservation efforts with the goal of improving deep learning models in the life sciences in a data-centric manner.

## 5 Acknowledgements

We are deeply grateful to Noelia Ferruz, Kevin Yang, Ahir Pushpanath, and Phoebe Oldach for helpful feedback and fruitful discussions throughout the writing of this manuscript. We thank Alexandros Papadopolous in particular for engineering support. We also want to thank Glen-Oliver Gowers, Oliver Vince, Sybil Wong, Leif Christoffersen, Bupe Mwambingu, Nadine Greenhalgh, Emma Bolton, Marlon Clarke, Ineke Knot, Neem Patel, William Chow, Carla Greco, Saif Ur-Rehman, Gus Minto-Cowcher, Keith Kam, Richard De Napoli, Gavin Ayres, Lily Goodyer Sait, and Marcus Leung.

## Supplementary Information

### A Methods

#### A.1 Global Sampling & Knowledge Graph Construction

Environmental samples subjected to metagenomic sequencing were collected after receiving landowner’s permission and entering access-benefit-sharing agreements with the relevant local or national authority, following Nagoya protocol guidelines. All samples were sequenced with both long-read and short-read sequencing methods applied after extraction. Alongside sample collection, we captured consistent metadata collection that include chemical, physical, weather, and geological measurements.

**Figure 1:**
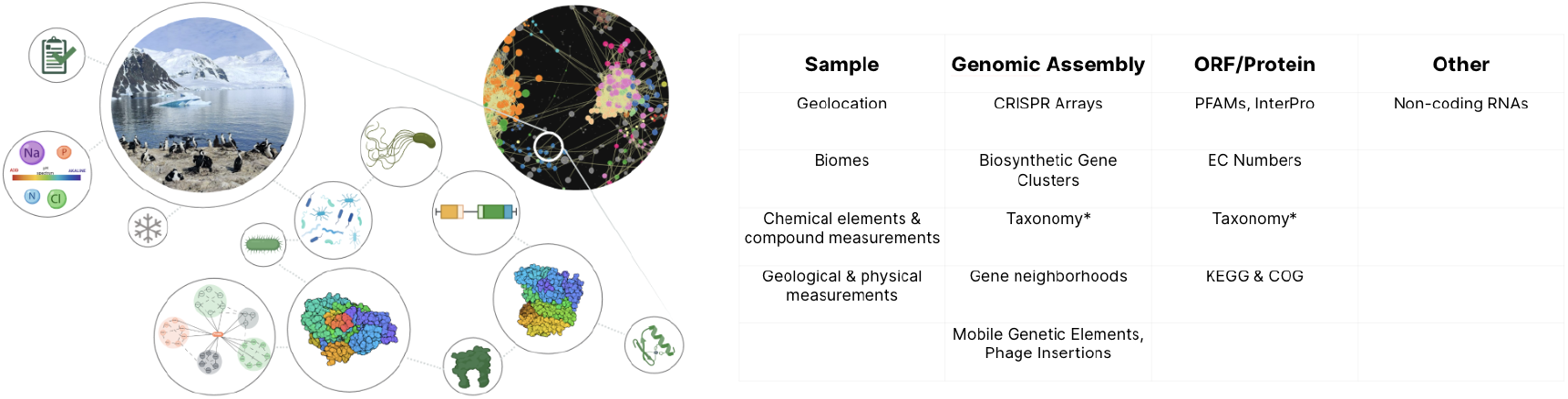
Visual representation of the data model for the Knowledge graph described in this study shown on the left. On the right we show a selection of information, measurements, and annotations associated with the entities in the graph. Taxonomies (*) are annotated both on the genomic assembly as well as the open reading frame (ORF) level.

We applied a custom assembly and annotation pipeline to the sequencing data which performs standard QC of sequencing reads and joint assembly of short and long reads to optimise for both low error rate and high assembly length. Open reading frames and non-coding RNAs are annotated on the genomic assemblies, along with CRISPR arrays, biosynthetic gene clusters, gene neighborhoods, mobile genetic elements, and phage integration events. The open reading frames were translated into amino acid sequences which were subjected to comprehensive *in silico* annotations, including PFAM [1], KEGG [2], COG [3], InterPro [4], and EC Numbers [6]. Functional annotations were performed with custom Hidden-Markov-Model-based and Deep Learning based models. Multiple custom taxonomic annotation methods were performed both on the ORF and the genomic assembly level. For an overview of these annotations and how they relate to the data model of the knowledge graph, see Figure 1.

For knowledge graph construction, we ingested all measurements, annotations, and information into a neo4j graph database.

#### A.2 Clustering

To assess the redundancy of protein sequence data across the Uniprot [1], [2], Mgnify [3], and Basecamp databases, we apply a hierarchical clustering strategy utilizing MMSeqs2 [4]. We initially cluster the Mgnify and Basecamp databases to a sequence identity threshold of 90%. Subsequently, we further refine the clustering of these databases down to 50% and 10% sequence identity thresholds. This stepwise reduction approach was chosen for its computational efficiency and reduced time consumption.

For the Uniprot database, we leverage the existing clustered datasets Uniref100, Uniref90, and Uniref50. These datasets provide a basis for our analysis, from which we identify the number of clusters at each identity threshold. Further, we utilize the Uniref50 clustered dataset to further cluster sequences down to a 10% identity threshold. This was achieved by adhering to the same clustering protocol used for the other datasets, which involves clustering sequences based on a 10% sequence identity and an 80% overlap with the longest sequence in the cluster.

#### A.3 Database Reduction For Efficient Search

To facilitate efficient searching within the environmental databases BRD v.2023.10 and Mgnify v.2022.05, we employed a database reduction strategy. This involved using the Linclust algorithm in Mmseqs2 to cluster both databases with a minimum sequence identity of 50% (–min-seq-id 0.5). This approach effectively reduced the combined database size to 239GB. After selecting cluster representatives, this process resulted in a total of 1.01 billion sequences used for the MSA supplemented flavour of AlphaFold2.

#### A.4 Template Search

AlphaFold2 [5] employs HHsearch [6] to scan a clustered version of the PDB (PDB70) for identifying the top 20 ranked templates. To maintain consistency with the original AlphaFold submissions for CASP15 targets, we configured the template search cutoff date to January 1, 2022. This setup was crucial to avoid any influence from newly deposited targets that might affect the predictions when using BRD.

#### A.5 Running BaseFold for Structural Comparison

To evaluate the full impact of the BRD on AlphaFold2 performance, we utilized AlphaFold’s default settings changing only the environmental database and template search date when computing the BaseFold structures. This approach aimed to directly compare the structural predictions under standard conditions. The CASP15 AlphaFold2 structures were downloaded from the AlphaFold2 Github repository and the AlphaFold2 structures for the CAMEO targets were downloaded directly from the CAMEO website. All BaseFold model inference was run on 8 NVIDIA A100 Tensor Core GPUs with 80GB of memory.

#### A.6 Molecular Docking

In preparation for docking, compound SAH was supplemented with Gasteiger charges followed by the addition of non polar hydrogen atoms [7]. Docking was performed using the default setting of AutoDock Vina [8] with a random seed of 42 and exhaustiveness set to 32. The box was defined using USCF Chimera [9] based on the crystal structure MFnG (PDB:7UX8). The defined dimensions of the box were 9.64 × 14.86 × 7.69 with a grid spacing of 1 Å, centered at coordinates x = 21.83, y = 38.56, z = 14.30 to maximize the precision of the substrate positioning within the active site. In the docking process, both the protein and ligands are treated as rigid entities. Results with a positional root-mean-square deviation (RMSD) below 1.0 Å were grouped, with each cluster represented by the most favorable binding free energy. The pose with the lowest binding affinity was then selected and aligned with the receptor structure for further analysis.

